# Striatal modulation supports policy-specific reinforcement and not action selection

**DOI:** 10.1101/2024.10.02.616336

**Authors:** A.T. Hodge, E.A. Yttri

## Abstract

Two contrasting models dominate our understanding of basal ganglia function: action selection and reinforcement learning. Prolonged, indiscriminate stimulation of direct and indirect pathway striatal neurons produces effects consistent with the action selection; however this approach ignores the transient, movement-specific dynamics that characterize these cells. To determine how striatal subpopulations contribute to mouse behavior, we applied brief closed-loop optogenetic stimulation to modulate ongoing activity in a manner that directly dissociates the contrasting models: upon the detection of locomotor arrest or leftward turns. While action selection models predict that increased direct pathway stimulation should induce locomotion and turning contralaterally to the side of stimulation, selective stimulation biased behavioral policies towards more frequent locomotor arrest and leftward turns, regardless of the side of stimulation. Indirect pathway stimulation had the opposite effect. Behavior followed the policy associated with the change in striatal activity, providing a mechanism to enable the reinforcement a wide range of behavioral features to shape performance.

## Introduction

Decision making is a dynamic process that underlies every action and its performance. The basal ganglia have been suggested to play an integral role in two facets of decision making: action selection and reinforcement learning (Amemori et al., 2011, 2018; Graybiel, 1995, 2005; Kravitz et al., 2010; Tai et al., 2012; Tecuapetla et al., 2014; Yin et al., 2005; Yttri & Dudman, 2016). Central to these proposed roles are the two populations that comprise the majority of the striatum, direct and indirect pathway spiny projection neurons - dSPNs and iSPNs, respectively (Balleine et al., 2009; Reig & Silberberg, 2014). dSPNs and iSPNs are thought to work in opposition to each other, mutually inhibiting each other and driving activity of the output nucleus of the basal ganglia in opposing directions (Burke et al., 2017; Tecuapetla et al., 2016). Some work has suggested that dSPNs select the action to be performed while iSPNs suppress unwanted actions (Kravitz et al., 2010; Tai et al., 2012; Tecuapetla et al., 2014). This premise of promoting and gating actions has been observed in to turning and orienting movements. dSPN stimulation promotes orienting contralateral to the stimulated hemisphere while iSPN stimulation induces ipsilateral turning, presumably due to inhibition of contralaterally oriented movements (Kravitz et al., 2010; Tai et al., 2012; Tecuapetla et al., 2014). Similarly, prolonged bilateral stimulation of dSPNs or iSPNs will induce or suppress locomotion. These effects are also consistent with the more generalized version of action selection, wherein dSPN and iSPN activity serve as ‘go’ and ‘no-go”signals, releasing or suppressing movement. In this framework, turning arises from the motoric go or stop being applied only unilaterally (Lauwereyns et al., 2002).

However, much of this work employs techniques that do not reflect the innate dynamics of these neurons. SPNs are mostly inactive, except transiently around the execution of a behavior. Continuous stimulation may require several seconds to elicit a behavioral response (Chang et al., 2008; Kravitz et al., 2010; Tecuapetla et al., 2014). Further, iSPNs and dSPNs are coactive during actions (Cui et al., 2013), complicating assertions concerning their respective stop and go roles, although some action selection models that take this observation into account (Bariselli et al., 2019). Finally, blocking the ultimate output of SPNs does not induce an akinetic state or lead to errors in action selection. Instead, inactivation or lesions of basal ganglia output nuclei only produce deficits in kinematic control and motor learning (Desmurget & Turner, 2010; Horak & Anderson, 1984; Kato & Kimura, 1992).

Alternative models of SPN function focus on their role in reinforcement learning, particularly surrounding the quality of performance. In these models, SPN activity adjusts features of a behavioral policy, e.g. vigor or speed, rather than selecting the policy outright (Shadmehr & Krakauer, 2008). Microstimulation of SPNs during successful associative learning trials increases future success rates while microstimulation during unsuccessful trials decreases success rates (Williams & Eskandar, 2006). Further, SPN firing rate directly correlates with learning rate during associative learning tasks. Pathway-specific stimulation in conjunction with lever presses leads to a progressive increase (dSPN) or decrease (iSPN) in the probability of future lever presses, and this bias persists after the cessation of stimulation (Kravitz et al., 2012). The persistence of these effects after the cessation of stimulation indicates that stimulation was modulating a gradual shift in behavioral policy rather than directly selecting a policy. Contrary to the “go” model of dSPN function, dSPN stimulation applied during slow movements increased the probability of slow movements in the future. iSPN stimulation had the opposite effect, demonstrating that there is not a direct mapping from dSPN activation to “move more” or iSPN activation to “move less.” (Yttri & Dudman, 2016). However, the extent to which these findings can be generalized to other contexts, particularly in the absence of a trained task, remains unclear.

How might we reconcile these hypotheses? Action selection models characterize SPNs as selecting specific actions. Regardless of when stimulation occurs, increased dSPN activity should have a fixed effect: the selection of a behavioral policy, e.g. “go/no-go”. The reinforcement model also predicts a fixed effect, but not with respect to specific actions. In this case, increased activity should bias the performance of behavior according to the context in which it occurs. With this in mind, we performed a series of experiments designed to directly dissociate these hypotheses and determine the computational contribution of striatal neurons to the execution of actions. We paired the detection of stopping or left turns with bilateral or unilateral optogenetic stimulation of dSPNs or iSPNs. dSPN stimulation upon stop detection led to extended periods of stopping and an increased likelihood of stopping, while iSPN stimulation upon stop detection led to large increases in locomotion. Importantly, speed during locomotor bouts was unaffected. Similarly, stimulation paired with left turn detection increased (dSPN) or decreased (iSPN) the probability of future left turns. This effect was consistent regardless of whether stimulation was delivered in the left hemisphere (ipsilateral), the right hemisphere (contralateral), or bilaterally. These effects were specific to the behavioral policy that increased SPN activity and demonstrate that modulation of striatal activity serves to reinforce features of behavioral policies in a more nuanced fashion than “go/no-go” action selection.

## Results

To determine the role of striatal activity in the selection or performance of actions, we placed mice in an open field and delivered bilateral stimulation at pseudorandom intervals. We stimulated dorsal striatum in animals expressing channelrhodopsin (ChR2) in dorsal striatal dSPNs or iSPNs (Drd1a-cre or Adora2a x ai32, Supplemental Figure 2A). Neurons in this somewhat medial part of striatum receive considerable input from forelimb motor cortex (Hooks et al., 2018; Yttri & Dudman, 2016) and demonstrate robust kinematic tuning during a variety of movements (Fobbs et al., 2020; Panigrahi et al., 2015).

As has been previously shown, prolonged (continuous, unpulsed stimulation for 30s) dSPN stimulation induced locomotion after several seconds, demonstrated as an increase in the average speed (Figure 1A, 300ms *p=0*.*992*, 1s *p<0*.*001*, 2s *p<0*.*001*). Conversely, prolonged open-loop iSPN stimulation decreased average speed (Figure 1A, 300ms *p=0*.*056*, 1s *p<0*.*001*, 2s *p<0*.*001*). These observations - the apparent facilitation or suppression of locomotion - constitute some of the strongest experimental evidence in support of striatum’s direct role in selecting what action to perform. However, brief (16 Hz for up to 450ms; see Methods) open-loop stimulation applied in the same pseudorandom manner did not induce locomotor changes (Figure 1B, dSPN 300ms *p=0*.*94*, 1s *p=0*.*59*, 2s *p=0*.*196*; iSPN 300ms *p=0*.*665*, 1s *p=0*.*854*, 2s *p=0*.*165*). The absence of an effect after only a short period of stimulation is consistent with the extended delay to effect observed for prolonged stimulation. However, near immediate changes in activity in downstream nuclei have been reported (Freeze et al., 2013; Roseberry et al., 2016), leaving open the source of this difference in effect.

**Figure 1:**
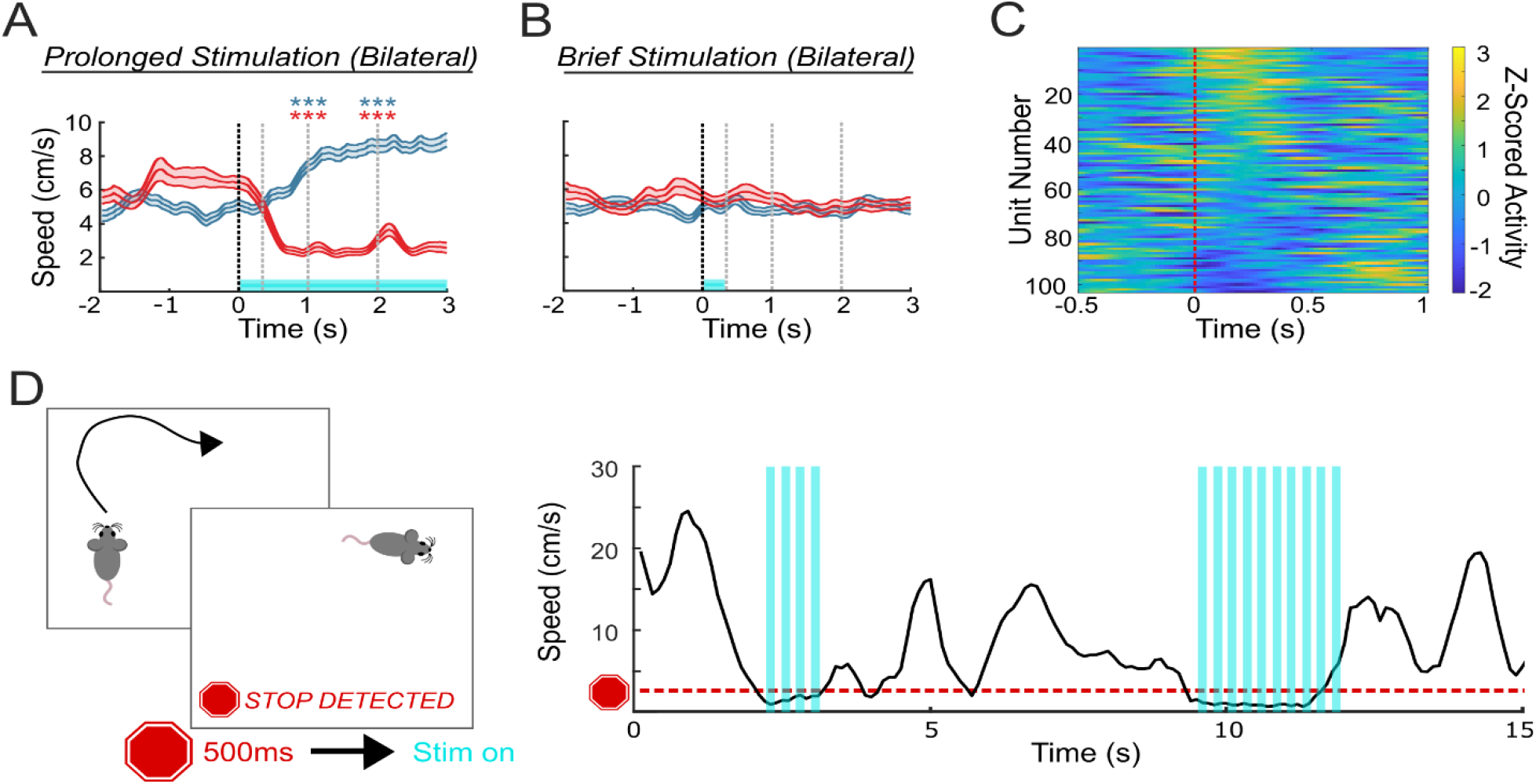
Prolonged and brief SPN stimulation produce distinct behavioral effects. **A)** While prolonged open-loop stimulation is sufficient to change locomotor speed, **B)** brief open-loop stimulation is not. Speed from animals receiving dSPN stimulation (blue) and from animals receiving iSPN stimulation (red). Cyan shading represents presence of stimulation. ^***^p<0.001, two sample t-test. **C)** Z-scored activity from putative SPNs in dorsal striatum aligned to stop detection. **D)** Schematic of stop-triggered stimulation design (left) and an example of closed-loop stimulation paired with stop detection (right). Using control data as an example, stimulation (cyan shading) was presented when locomotor speed decreased below 2cm/s for at least 500ms.

The shorter period of stimulation provides the opportunity for more accurate credit assignment relating to a particular behavioral phenomenon. Although neither approach is specifically designed to mimic specific recorded patterns of SPN activity, brief stimulation limits the increases in SPN activity to the brief periods of transitory activity observed *in vivo* during locomotion and stopping (Figure 1C, Jurado-Parras et al., 2020; Rueda-Orozco & Robbe, 2015).

### Selective dSPN stimulation can decrease locomotor output, while iSPN stimulation increases locomotor output

To dissociate the potential role in reinforcement from the selection of behavior, we applied brief dSPN or iSPN stimulation when locomotor arrest (henceforth, ‘stop’) was detected, e.g. when locomotor speed fell below 2cm/s for 500ms (Figure 1D). Although pro-movement models suggest that dSPN stimulation should increase locomotor speed, brief dSPN stimulation following stopping instead increased stop probability. We note that brief stimulation following stop detection did not select a specific locomotor outcome. For instance, although selective dSPN stimulation biased mice towards prolonged stopping bouts (locomotor bout initiation decreased in the second following stimulation *p=0*.*017*), we regularly observed the initiation of locomotor bouts during dSPN stimulation (Figure 2A). While speed decreased following dSPN stimulation (Figure 2B, dSPN stim mean peak speed = 3.92 cm/s, control mean peak speed = 4.07 cm/s, *p<0*.*001*), this was more consistent with an increase in stopping than a general decrease in speed. The interval between detected stopping bouts decreased (Fig 2C median inter-stop interval: control=1.60s, stim=1.39s; *p=0*.*01*) and the duration of the stop bouts increased (Figure 2D median stop duration: control=2.56s, stim=3.29s, *p=0*.*015*). However, dSPN stimulation did not induce a change in peak speed during locomotor bouts (Figure 2E median dSPN control=15.4cm/s, median dSPN stim=15.6cm/s, *p=0*.*63*). These results are consistent with increased dSPN activity following stopping reinforcing stopping bouts rather than selecting locomotor speed.

**Figure 2:**
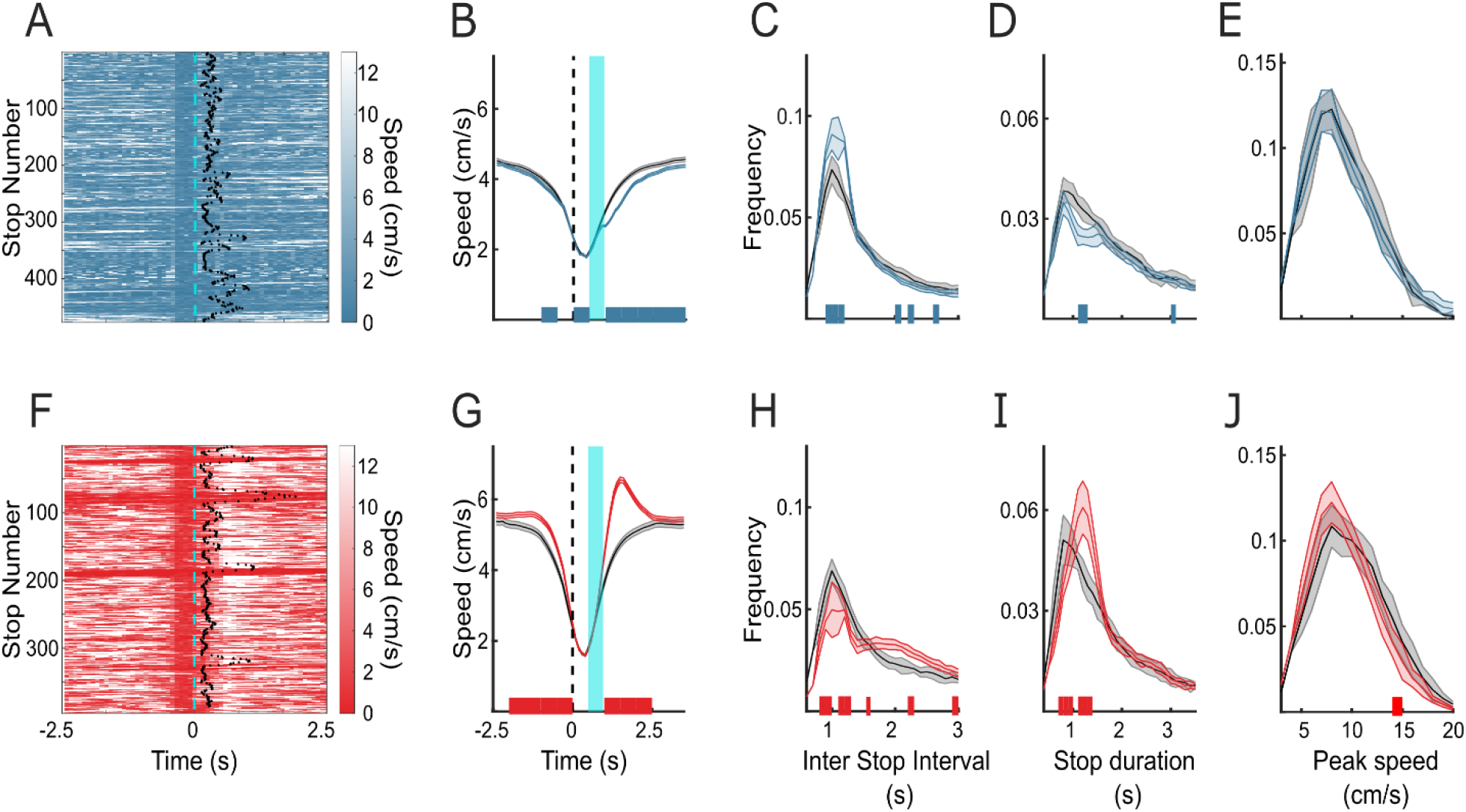
Stimulation of SPNs triggered by stop detection reinforces stopping. **A)** Speed aligned to the administration of dSPN stimulation (dashed cyan line) in one example session. Black dots indicate the first point after each stop where speed increased above the 2cm/s threshold. **B)** locomotor speed aligned to stop detection in dSPN stimulation (blue) and control (black) sessions. Stimulation period denoted by cyan shading. Significant differences (p<0.001) between stimulation and control are denoted by blue ticks on the x axis. **C)** distribution of inter-stop intervals in dSPN stimulation (blue) and control (black) sessions. Significant differences (p<0.05) between stimulation and control are denoted by blue ticks on the x axis. **D)** distribution of stop durations in dSPN stimulation (blue) and control (black) sessions. Significant differences (p<0.05) between stimulation and control are denoted by blue ticks on the x axis. **E)** distribution of peak speeds during locomotor bouts in dSPN stimulation (blue) and control (black) sessions. Significant differences (p<0.05) between stimulation and control are denoted by blue ticks on the x axis. **F-J)** as **A-D)**, but for iSPN stimulation (red) sessions.

Conversely, mice receiving iSPN stimulation were less prone to locomotor arrests (Figure 2F), with locomotor bout initiation increasing in the second following stimulation (*p=0*.*039*). Speed also transiently increased following iSPN stimulation (Figure 2G, iSPN stim mean peak speed = 5.45 cm/s, control mean peak speed = 4.99 cm/s, *p<0*.*001*) and locomotor bouts between stops became longer (Figure 2H median inter-stop interval: control=1.69s, iSPN stim=2.00s; *p=0*.*003*). iSPN stimulation induced a non-significant trend towards shorter stop durations (Figure 2I control=2.52s, stim=2.21s; *p=0*.*09*), with the strongest effect just after initial stimulation concluded. iSPN stimulation did not influence peak locomotor bout speed (Figure 2J median iSPN control=17.8cm/s, median iSPN stim=16.3cm/s, *p=0*.*25*). These results indicate that increases in iSPN activity following stopping negatively reinforced stopping without directly selecting locomotor speed. This contradicts current action selection and go/stop models of striatal function. Instead, they demonstrate that increases in SPN activity corresponding to behavioral policy (Rueda-Orozco & Robbe, 2015) produce a policy-specific reinforcement of the selected behavior. In this case, stimulation in conjunction with near-zero locomotor speed led to either an increase (dSPN) or a decrease (iSPN) in future stopping.

We recognize that reinforcement mechanisms can lead to place preference (CPP), creating the potential that some of these effects might be due to the motivation to persist within an area. To dissociate locomotor arrest from a spatial preference for a given area, we divided the arena into quadrants and determined the percent of time spent within the most-preferred through least-preferred quadrant for that session. There was no difference in the number of entries into the most preferred quadrant of the open field or in the number of entries into the least preferred quadrant of the open field (Figure 3C dSPN most preferred *p=0*.*472*, least preferred *p=0*.*555*, two sample t-test; Figure 3H iSPN most preferred *p=0*.*864*, least preferred *p=0*.*602*, two sample t-test). We also found no change in the occupancy of the preference-ranked quadrants resulting from dSPN or iSPN stimulation (Figure 3D dSPN most preferred *p=0*.*887*, least preferred *p=0*.*636*, two sample t-test; Figure 3I iSPN most preferred *p=0*.*336*, least preferred *p=0*.*164*, two sample t-test). Additionally, while we observed a change in the total number of stops that occurred as a result of dSPN or iSPN stimulation above, the proportion of stops in each quadrant did not change (Figure 3E dSPN most preferred *p=0*.*766*, least preferred *p=0*.*839*, two sample t-test; Figure 3J iSPN most preferred *p=0*.*335*, least preferred *p=0*.*715*, two sample t-test). These results indicate that SPN stimulation did not induce off-target CPP, instead selectively reinforcing the policy feature that triggered stimulation.

**Figure 3:**
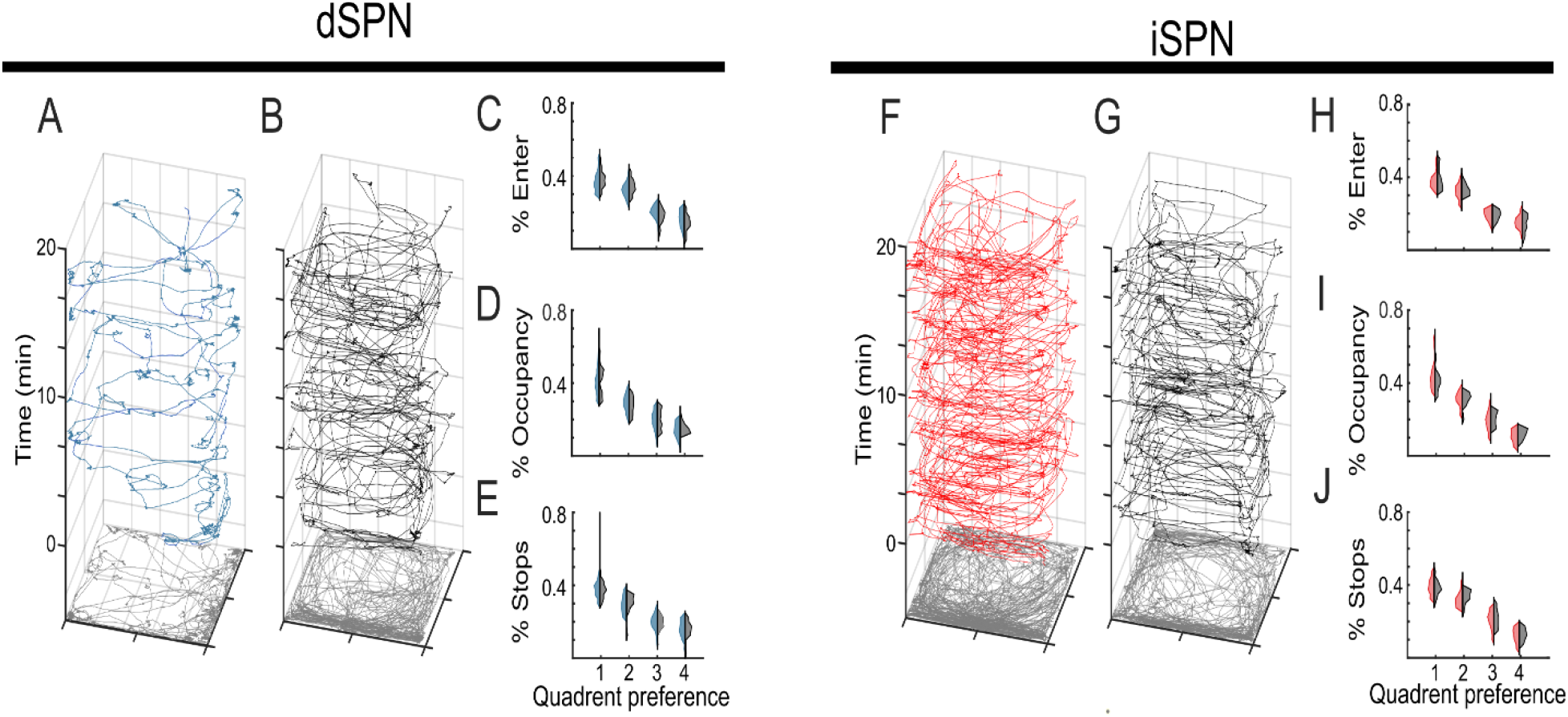
Stop-triggered SPN stimulation does not produce off-target reinforcement. Example locomotor trajectories throughout 20-minute **A)** dSPN stim (blue) and **B)** dSPN sham session (black). Gray trace below indicates mouse position collapsed across time. Data are from the same mouse. Proportion of **C)** entrances, **D)** occupancy time, **E)** and stop events for each of the four quadrants. In each violin plot, blue indicates data from stimulation sessions, black indicates data from sham sessions. Ranked preference for the four quadrants was determined independently for each session. **F-J)** Same as above, but for iSPN stimulation. No significant differences were found (two sample t-test)

### SPN modulation reinforces turning, regardless of side of stimulation

In the open field, bilateral SPN stimulation reinforces a specific feature of a behavioral policy (stop frequency), arguing that SPN activity reinforces behavior rather than driving actions. It has been observed that prolonged unilateral open-loop dSPN stimulation compels the execution of wide turns contralateral to the stimulated side, while prolonged unilateral iSPN stimulation induces tight ipsilateral turns. This orienting behavior has been interpreted as the selection or suppression of direction-specific action channels (Vich et al., 2022)that then generate turns. However, we found similar brief SPN activity in the dorsal striatum in the left hemisphere aligned to either left or right turns (Figure 4A). This suggests that the activity of individual SPNs do not select turn direction.

**Figure 4:**
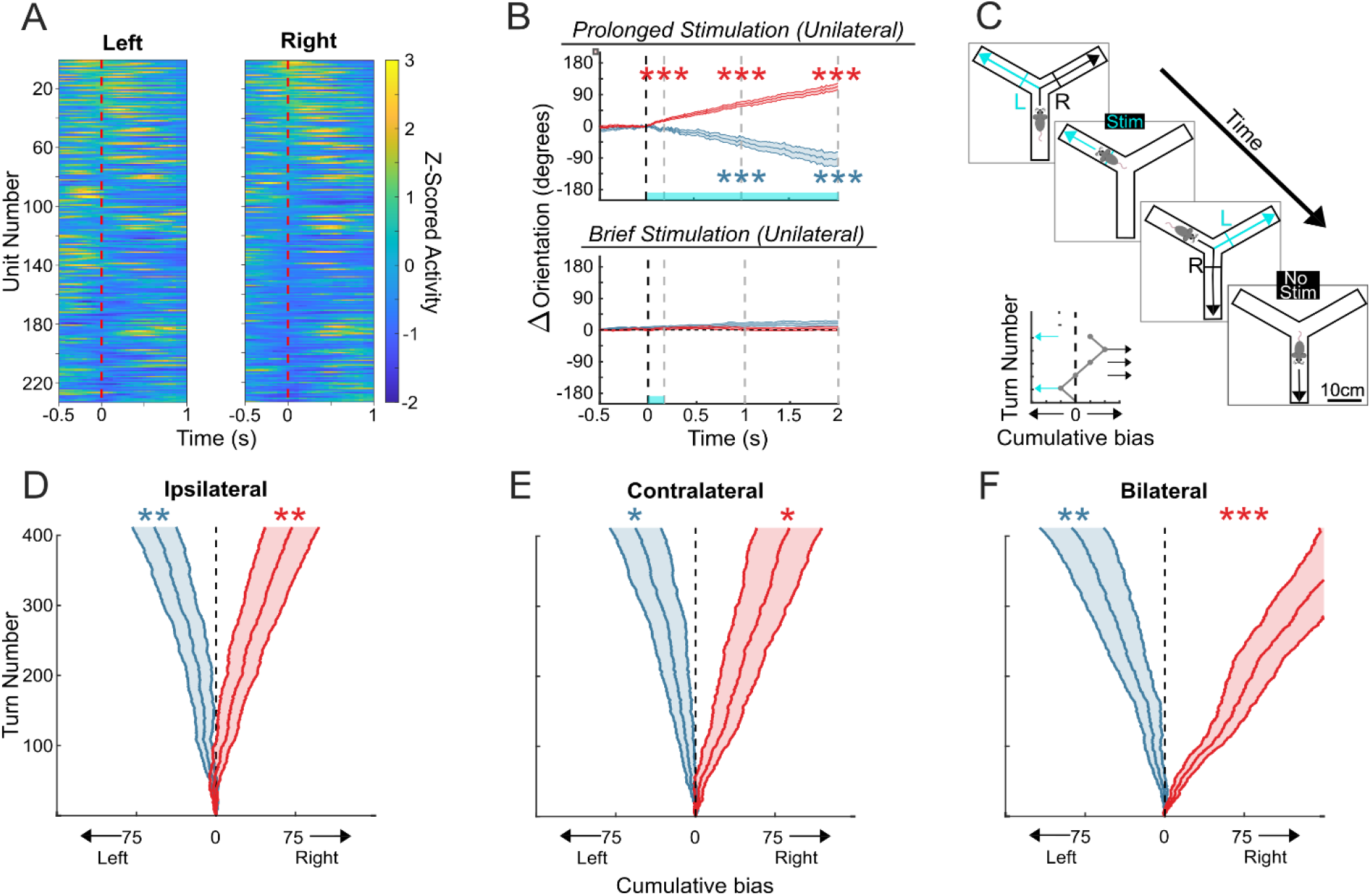
SPN stimulation reinforces turn direction. **A)** Z-scored activity of putative SPNs in dorsal striatum aligned to left turn detection (left side) or right turn detection (right side) in a Y-maze. SPNs were similarly active during turns in either direction. **B)** Effect of prolonged (top) and brief stimulation (bottom) on orientation heading in the open field. Positive values indicate ipsilateral turning. ^***^ p<0.001, 2-sample t-test for a given timepoint (300ms, 1s, or 2s) compared to pre-stimulation baseline. **C)** Diagram of the Y-maze experimental paradigm. Left turn detection triggers brief optogenetic stimulation of dSPNs or iSPNs, regardless of starting hall. Inset is an example of a bias plot covering 10 turns within one session. A higher proportion of left turns will cause the line to move left. **D-F)** Mean cumulative turn bias, e.g. the difference between Left and Right turn counts after a given number of total turns, for sessions in which stimulation occurred **D)** ipsilaterally, **E)** contralaterally, or **F)** bilaterally relative to the turn direction (left). shaded regions represent mean +/-SEM. ^*^p<0.05, ^**^p<0.01, ^***^p<0.001, one-sample t-test.

When we applied open-loop, prolonged unilateral stimulation, we also observed the expected contralateral or ipsilateral turning (Figure 4B top dSPN 300ms *p=0*.*061*, 1s *p<0*.*001*, 2s *p<0*.*001*; iSPN 300ms *p<0*.*001*, 1s *p<0*.*001*, 2s *p<0*.*001*). However, brief stimulation failed to elicit a turning response in the same mice (Figure 4B bottom dSPN 300ms *p=0*.*56*, 1s *p=0*.*12*, 2s *p=0*.*07*; iSPN 300ms *p=0*.*24*, 1s *p=0*.*57*, 2s *p=0*.*5*).

We then constrained turning behavior to ensure that the only feature that differed between leftward and rightward turns was direction (Figure 4C). ‘Turning’ is a behavioral policy with a large range of potentially relevant features, making the clear and specific reinforcement of just one (i.e. direction) non-trivial (see Supplemental Figure 3 for discussion about the difficulties of defining what all may constitute a ‘left turn’). A Y-maze provides improved spatiotemporal resolution and consistency for detecting the occurrence of turning decisions (Friedman et al., 2015, 2017). To motivate turning, entry into any hall carried a 15% chance of liquid reward delivery, regardless of turn direction. Rewards were delivered via lick port located at the end of each arm of the maze.

We paired brief optogenetic stimulation of dSPNs or iSPNs with the detection of the execution left turns in a Y-maze (Figure 4C, cyan lines). Stimulation was administered in either the left hemisphere (ipsilateral), the right hemisphere (contralateral), or bilaterally. Regardless of the side administered, dSPN stimulation significantly increased the probability of performing left turns (Figure 4D-F, one-sample t-test on difference in cumulative bias: ipsilateral stimulation *p<0*.*01*, contralateral stimulation *p=0*.*03*, bilateral stimulation *p<0*.*01*). Conversely, iSPN stimulation significantly decreased left turns – again, regardless of the stimulation side (Figure 4D-F, one-sample t-test, ipsilateral *p=0*.*01*, contralateral *p=0*.*01*, bilateral *p<0*.*001*, respectively). The difference between left and right turn counts increased at a steady rate within individual sessions, suggesting that increased SPN activity after turn detection was contributing to a progressive reinforcement of one specific direction (i.e. left). These changes in performance were specific to the choice of left or right turn detection; we found no effect of stimulation on hall preference (Supplemental Figure 4, all comparisons *p<0*.*05* 2-way ANOVA with Bonferroni multiple corrections) or locomotor speed (dSPN Ipsilateral *p=0*.*75*, Contralateral *p=0*.*94*, Bilateral *p=0*.*91*; iSPN Ipsilateral *p=0*.*17*, Contralateral *p=0*.*47*, Bilateral *p=0*.*21*, two sample t-test).

To control for the effects of the random reward, including any potential overlap in eligibility trace between stimulation and reward, naïve mice were placed in the Y-maze without reward. Although the rate of turn execution was greatly reduced in the absence of reward, the effect of SPN stimulation was preserved. Left hemisphere dSPN stimulation biased mice to perform more ipsilateral turns (Figure 5A, *p=0*.*029*), while iSPN stimulation in the left hemisphere produced a contralateral turn bias (Figure 5A, *p<0*.*0003*). Importantly, there was also no difference in the average slope of cumulative bias values in rewarded and unrewarded contexts, either at the unrewarded session trial cutoff (Figure 5B, Turn number=120 turns, dSPN *p=0*.*94*; iSPN *p=0*.*83*). or at the respective cutoffs (rewarded = 412 turns, unrewarded = 120; dSPN *p=0*.*28*; iSPN *p=0*.*72*). These results demonstrate that increased SPN activity reinforces performance towards or away from a paired context, such as the direction of a turn, including contexts in which an animal does not have an explicit external motivation for performing an action.

**Figure 5:**
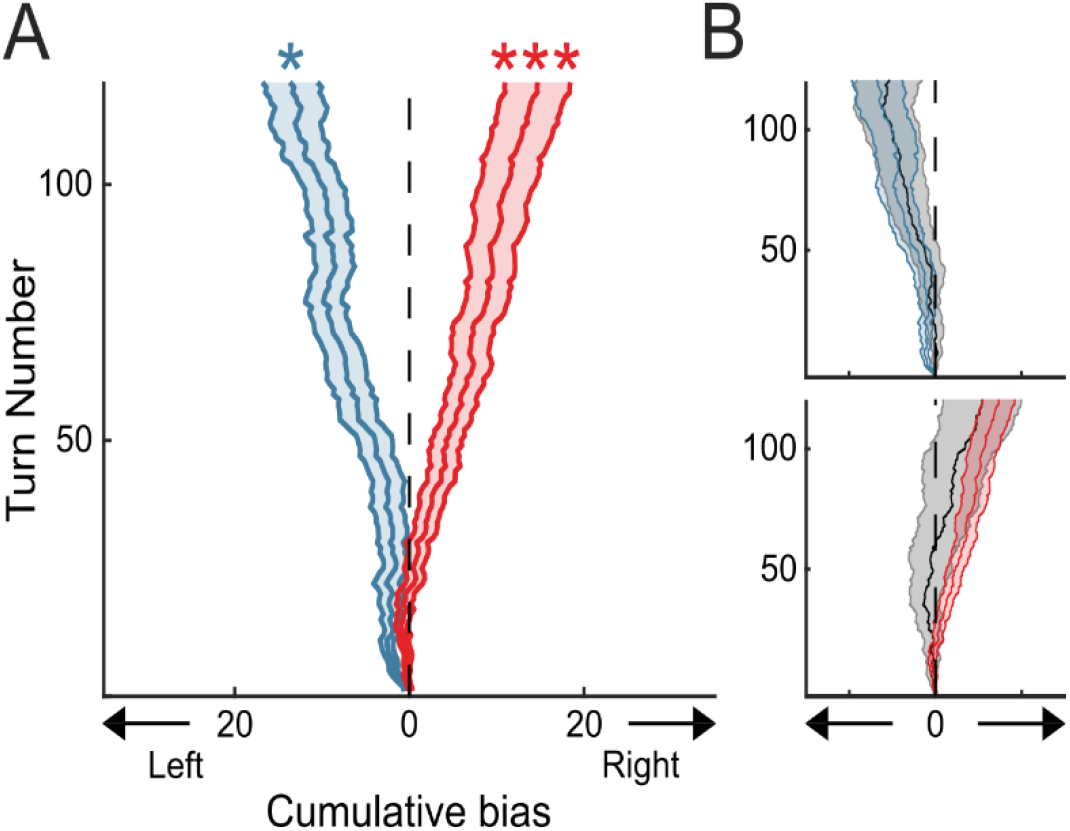
SPN stimulation reinforces direction bias in the absence of reward. **A)** Cumulative turn bias in unrewarded sessions where dSPN (blue) or iSPN (red) activation occurred ipsilaterally to the turn direction. ^*^p<0.05, ^***^p<.001; one-sample t-test for final within-animal baseline subtracted bias values. For comparison, effects of **B)** dSPN or iSPN stimulation from unrewarded and rewarded (gray) sessions are shown together through the first 120 trials.

## Discussion

Two theories dominate our understanding of the contributions of the basal ganglia in motoric decision-making: action selection and reinforcement. While related, these functions differ in several key ways. Here, we provide evidence that striatal SPNs do not act to select a given behavior, but rather to reinforce particular aspects of it with great specificity. These reinforcing effects may extend to a range of behavioral dynamics and specific kinematic elements of behaviors (such as orientation during locomotion). However, these findings stand in contrast to prevailing notions that dSPN activity is pro-movement and iSPN activity is anti-movement.

The canonical models of basal ganglia function were created to describe the hypo- and hyperkinetic shifts in performance associated with diseases of the basal ganglia (Georgopoulos et al., 1983; Albin et al., 1989). Newer theories pivoted from modulating vigor to the promotion or suppression of movement or individual actions (Lauwereyns et al., 2002; Mink, 1996). Interestingly, the disease states that served as the foundation for these models do not demonstrate deficits in action selection. Parkinson’s disease patients do not select inappropriate actions, but rather, demonstrate impaired adaptive gain control of various movement types (Shadmehr & Krakauer, 2008). Similarly, lesions of the output nuclei of the basal ganglia yield only learning and kinematic deficits, leaving even complex action selection intact (Turner et al., 2007).

Our findings demonstrate that the striatum serves as a nexus for policy-based reinforcement. This is consistent with the effects of selective electrical stimulation of the caudate in primates. Learning a joystick task was facilitated when stimulation occurred in conjunction with the desired performance, while the same stimulation paired with undesirable performance drove learning in the opposite direction (Williams & Eskandar, 2006). When not paired with performance, stimulation had no effect. Kravitz (2012) also demonstrated the importance of context. showed that prolonged stimulation of SPNs following a specific behavior (lever press) reinforced that feature in a manner that was similar to the results we’ve described here. Our data are consistent with the hypothesis that the enacted motor command is formulated and executed upstream of the striatum. In such a model, these motor plans are passed through the basal ganglia, where their kinematic performance is adjusted according to its context (Dudman & Krakauer, 2016). Proceeding downstream from the basal ganglia, 60-70% of mammalian basal ganglia efferent projections consist of midbrain and brainstem nuclei (McElvain et al., 2021; Parent et al., 1983; Parent & De Bellefeuille, 1983), including those with well documented motor function like the superior colliculus and mesencephalic locomotor region (MLR). Here, the basal ganglia projections converge with the same cortical neurons that project to the striatum (Tervo et al., 2016), providing a means to connect the computations in striatum with the descending motor commands.

How then might we reconcile the effects of prolonged, open-loop stimulation with the functional architecture of the brain? The pro-movement effects of prolonged stimulation may be the result of cumulative or off-target effects. Over time, the continuous, maximal activation of the striatum may activate or disinhibit downstream premotor effectors far beyond the targeted neurons, notably the superior colliculus and MLR. Manipulation of either of these areas induces robust changes in behavior that are akin to prolonged SPN stimulation. The generation of orienting turning movements, either of the body or the eyes, is a well-known consequence of superior colliculus stimulation (Basso & May, 2017; Zahler et al., 2021), and activation of the MLR has similarly been shown to drive locomotion (Caggiano et al., 2018). Thus, while no causal perturbation can assert complete physiological fidelity, there are significant considerations to keep in mind when comparing ‘scalpel or sledgehammer’ approaches.

We note that closed-loop stimulation occurred in the midst of the behavior of interest. This was also the case in our previous work in which stimulation occurred only after the detection of a forelimb reaching movement (Yttri and Dudman 2016). As one cannot assure accurate prediction of the impending motor performance before its onset, the application of behavior-specific stimulation must wait until the detection of that behavior is complete. Although this technical limitation may bias our findings towards learning effects on subsequent performance, we would reiterate that the identical pattern of stimulation delivered in open-loop does not affect the selection of action, nor does this brief, random stimulation induce even modest changes in speed upon stimulation onset. Critically, these experiments demonstrate effects that cannot be explained by action selection models.

## Acknowledgements

We thank Dr. Joshua Dudman for support in gathering pilot data for this study. We would like to acknowledge Mark Nicholas for providing surgical expertise. We also thank members of the Yttri lab, Drs Aryn Gittis, Daniel Leventhal, and Sandra Kuhlman for their insightful comments. This work was supported the Whitehall foundation and Kaufman foundation grants, as well as NIH CRCNS grant R01DA053014.

## Author contributions

ATH conceived the study, acquired, analyzed and interpreted the data, created the analysis software and drafted the manuscript. EAY conceived the study, analyzed and interpreted the data, and drafted the manuscript.

## Data Availability

Data and MATLAB code used to perform analyses/construct figures are available at https://github.com/athodge/Hodge_and_Yttri_2024/

## Methods

### Subjects

Adult (>p60) double transgenic drd1-cre x ai32 mice (for the expression of ChR2 fused to enhanced yellow fluorescent protein in cells expressing D1 dopamine receptors, 9 males, 7 females) or adora2a-cre x ai32 transgenic mice (for the expression of ChR2-EYFP in cells expressing Adora-2A adenosine receptors, 5 males, 8 females) were used, all on a C57BL/6J background. Mice were housed in temperature and humidity-controlled facilities on a reversed 12-hour light/dark cycle. All experiments were approved by Carnegie Mellon University Institutional Animal Care and Use Committee.

### Surgical procedures

All surgeries were performed under anesthesia (2% isoflurane gas anesthesia). Optic fibers were stereotaxically implanted in dorsal striatum (coordinates from bregma: anterior 0.5, lateral 1.5, and -1.5 to -1.75mm ventral to dura) though burr holes. This location within the striatum receives input from M1 (Sup Fig 2, see also Yttri and Dudman 2016 Extended Data Fig 1) and is sufficient to drive movement following prolonged stimulation. Optic fibers were secured with cyanoacrylate glue and dental acrylic (OrthoJet). 3D printed PLA head caps used to assist in animal tracking were attached during the same procedure. Mice received intraperitoneal carprofen injections (5mg/kg) before surgery and on the first day of recovery. Further injections were administered as needed. All procedures were performed under aseptic conditions. Mice were monitored throughout recovery and returned to home cages where they were housed individually or group housed singly through the use of grated aluminum cage dividers.

To verify anatomical location of the fiber within dorsal striatum, animals were anaesthetized and transcardially perfused with ice-cold phosphate buffer saline (pH 7.45) and ice-cold paraformaldehyde (4% pfa in pbs). Brains were placed in paraformaldehyde overnight and then moved to a 4% sucrose solution a day prior to sectioning. 100 um sections were mounted using a DAPI-stain mounting medium and imaged with a Zeiss Axioplan2 upright light/fluorescence microscope.

### Optogenetic stimulation

Light source (Prizmatix Optogenetics LED) power was calibrated to 1-2mW at fiber tip (0.66 NA, 200um diameter). For brief stimulation, 460nm light was delivered in 10ms pulses at 16 Hz for 450ms (Figures 1 through 3) or 310ms (Figures 4 and 5). Stimulation duration was shortened in the Y-maze in an attempt to limit stimulation to the duration of the turn. For prolonged stimulation, the same power was used, but with stimulation applied continuously for 30s.

### Animal tracking

For stop-triggered stimulation experiments, mouse position was sampled at 10Hz by an overhead camera that utilized a custom blob detection algorithm. In the other experiments, mice were fitted with two-hued markers in the open field or single-hued markers in the Y-maze (for improved maneuverability) and allowed to habituate to the weight (∼0.5 or 1g) for 24hrs. Animal position was captured via PixyCam2 (Charmed Labs) computer vision system in conjunction with colored markers attached to mouse headcaps. Orientation was recorded online via the relationship of the two hues of the position marker. The majority of our data were collected prior to the availability of more detailed behavioral extraction techniques, including B-SOiD (Hsu & Yttri, 2021). Anecdotally, mice did not change posture upon as a result of brief stimulation while stopped, nor was a stereotyped action such as a freezing fear response noted.

### Open-loop stimulation

Open-loop bilateral and unilateral stimulation in the open field was administered in manner similar to Kravitz et al., 2010. For prolonged stimulation sessions, a total of ten stimulations were administered with a 1-minute inter-stimulation intervals after a 1 to 10-minute acclimation period. Sessions lasted for 20 minutes and occurred at most once per day. For brief stimulation sessions, after a 10-minute acclimation period, a pseudo-random number of stimulations was administered with a mean inter-stimulation interval of 30s over the course of 10 minutes. For prolonged and brief unilateral stimulation sessions, stimulation was applied in the left hemisphere in 19 of 21 sessions, with the effects of the remaining right-side stimulation sessions flipped such that turn laterality relative to stimulation side was maintained. The same animals comprise the Y-maze and open-loop unilateral stimulation datasets. Different animals comprise the open field and open-loop bilateral stimulation datasets.

### Stop-triggered experiment

Following a 20-minute acclimation period to a 1m x 1m arena, 450ms pulsed stimulation was applied bilaterally whenever locomotor arrest (“stopping”) was detected. A stopped state was defined as a cumulative translation in space totaling less than 2cm in a 500ms period. 500ms sampling periods were consecutive and non-overlapping. This experimental period lasted for 20 minutes. As in all control sessions, optogenetic patch cables were attached to the animal but no stimulation was applied during this time. Before experiments began, mice had experienced the arena without stimulation for three, one-hour sessions.

### Y-maze experiment

Mice freely traversed a Y-maze with halls each measuring 20cm long, 45cm tall, and with an interior width of 5cm. Head position information was sent to an Arduino running custom code for turn detection and data acquisition. Turns were defined as having exited the currently occupied hall and proceeding 6.6cm down a different hall. This distance was chosen as it is greater than ∼75% of a body length (Chakraborty et al., 2017; Dunn & Bale, 2009). Thus, detection occurs when a mouse has fully committed to a turn but is still in the process of executing the turn. This threshold ensured that stimulation only occurred in conjunction with turns and not investigative pokes, e.g. vicarious trial and error behavior (Meunzinger & Gentry, 1931; Redish, 2016; Tolman, 1939, 1948). Turn direction was defined egocentrically from the animal’s perspective. Mice were water restricted for three days and acclimated to the Y-maze for several sessions prior to the collection of control or experimental data. Y-maze sessions lasted for 30 minutes and began within 30 seconds of placing the animal in the maze. The variable number of turns per sessions was addressed by setting a minimum turn threshold for inclusion of 15% of the maximum turn count across sessions, equating to 120 turns for unrewarded sessions or 12 turns for rewarded sessions. This inclusion threshold was set to maintain a similar turn rate and trial count when collapsing across sessions. Sessions that did not meet this threshold were excluded from all analyses. Turns within a session in excess of the threshold were discarded. In the unrewarded context, mice experienced at five control (no stimulation) and ipsilateral stimulation sessions.

The reward-naïve mice were then introduced to the rewarded context, which they experienced for at least three acclimation sessions before data was collected. Five sessions each of control (no stimulation), ipsilateral stim, contralateral stim, and bilateral stimulation conditions pseudo-randomly interleaved. Liquid reward (3mM acesulfame potassium, Ace-k) was delivered at lick ports located at the end of each hall. Every turn detection (hall entry) carried a 15% chance of actuating an audible solenoid (3-port HDI, The Lee Company, Essex, CT) associated with that hall.

### Electrophysiology

To verify that dorsal striatal neurons modulated their activity around the time of stimulation onset, we chronically implanted mice with Neuropixels electrodes (IMEC) at approximately 0.5 mm anterior, 1.8 mm lateral of bregma at a depth of at least 2.1 mm below the surface of the brain using aseptic conditions. Data acquisition was performed using OpenEphys hardware. Spiking activity was then band-pass filtered (300 Hz-3 kHz) and sorted offline using Kilosort3. Putative SPNs were conservatively identified as those well-isolated neurons having a mean firing rate under 6Hz. Spikes of isolated single units were counted within 20ms bins to generate the average response per unit aligned to behavior detection. The period encompassing one second before and after alignment was used for Z-score normalization.

### Statistics

All analyses were performed with MATLAB (MathWorks). Unless otherwise specified, parametric testing utilized one-sample or two sample t-tests and nonparametric testing for non-normal distributions utilized two sample Kolmogorov-Smirnov tests. In figures, results are presented as mean + S.E.M of each distribution unless otherwise noted. Mouse-to-mouse choice variability was controlled for in Y-maze contexts through within-animal baseline subtraction. Mean behavior from control sessions without stimulation from a given mouse was subtracted from that mouse’s stimulation sessions. Sessions were pooled within stimulation type and genotype following within-animal baseline subtraction.

We wish to be clear about our terminology involving reinforcement learning, including differences between operant conditioning and some parts of the machine learning fields. Within the field of operant conditioning, the term ‘positive’ or ‘negative’ refers to whether a *stimulus* is added or removed, respectively. Thus, ‘positive punishment’ is the reduction of the occurrence of a behavior through the addition of a stimulus – as was observed via iSPN stimulation - while ‘negative reinforcement’ is the increased occurrence of a behavior through the removal of a stimulus (see Kravitz et al., 2012 for review). To avoid confusion, we have avoided referring to valence of reinforcement or punishment, grouping everything under the broader concept of ‘reinforcement’. We also note that there should be no assumption of a hedonic or appetitive value as a result of stimulation, e.g. “liking.”

