## Supplemental Figures for "Striatal modulation supports policy-specific reinforcement and not action selection"

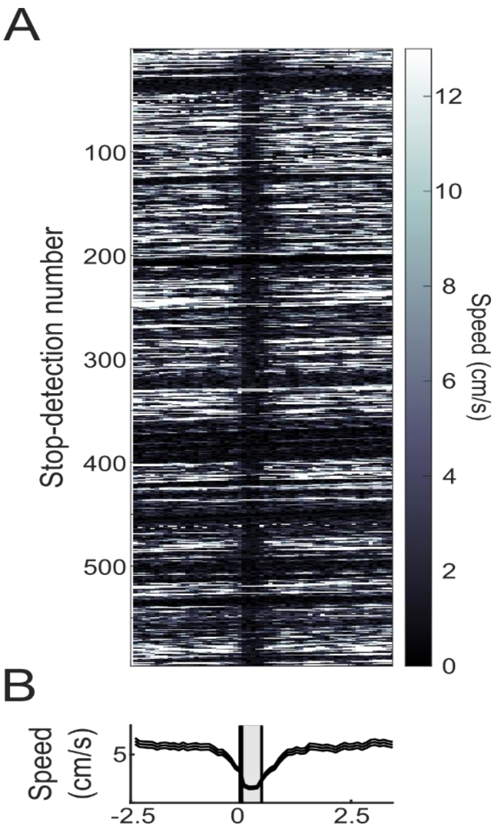


**Supplemental Figure 1: Performance of our stop detection method during a control session. A)** Each row represents one identified period of 500ms locomotor arrest. Prolonged stopping, e.g. consecutive 500ms periods of locomotor arrest, will be seen as stacks of horizontal black bars. **B)** Average speed aligned to stop detection

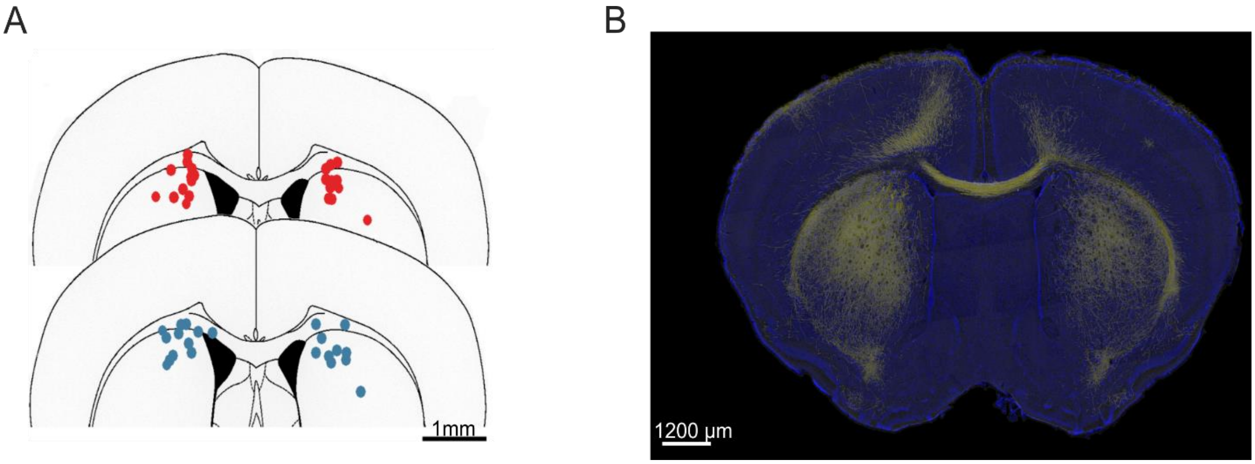


**Supplemental Figure 2: Motor cortex projects to dorsomedial and dorsolateral striatum.**
**A)** Diagram of fiber optic implant tip locations. Unless otherwise specified, all mice received bilateral fiber optic implants in the dorsal striatum. Red dots (Top) indicate the tip locations of all A2A-cre x Ai32 mice used in this study. Blue dots (bottom) indicate the tip locations of all D1-cre x Ai32 mice used in this study **B)** GFP injection into forelimb M1 demonstrates innervation across the mediolateral axis of dorsal striatum.  Image provided by Dr. Mac Hooks, University of Pittsburgh, from unpublished slides related to “Topographic precision in sensory and motor corticostriatal projections varies across cell type and cortical area” Nature Communications, 2018

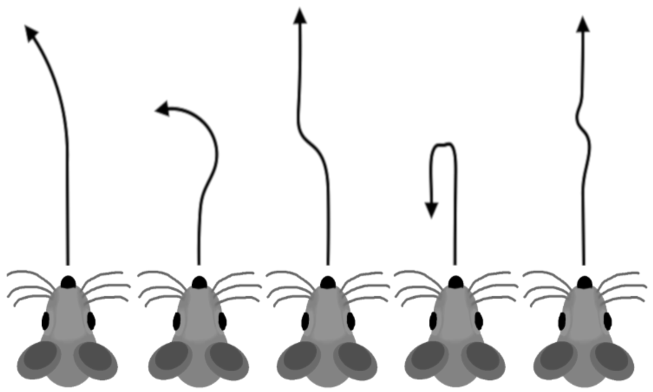

**Supplemental Figure 3: Variations and nuance in possible turn definitions.** While it is commonly agreed upon that a turn should constitute a change in heading over some distance, precise definitions and thus detection can be deceptively difficult. In an open field, the difference between a turn and a slanted forward trajectory is ambiguous. Likewise, it is unclear whether a trajectory with multiple deviations in heading is a single turn with multiple components or multiple small turns. It is also unclear whether large (>180 degrees) and small (<90 degrees) turns are distinct behaviors or variations in the execution of the same behavior.

Mazes, including Y-mazes and T-mazes, limit the possible range of actions that can be described as ‘Turns.’ Unlike the often-used T-maze, a Y-maze is radially symmetric, making the egocentric left/right decision identical regardless of the hall of origin. This feature reduces experimenter intervention, increasing trial counts and minimizing experimenter-introduced confounds.

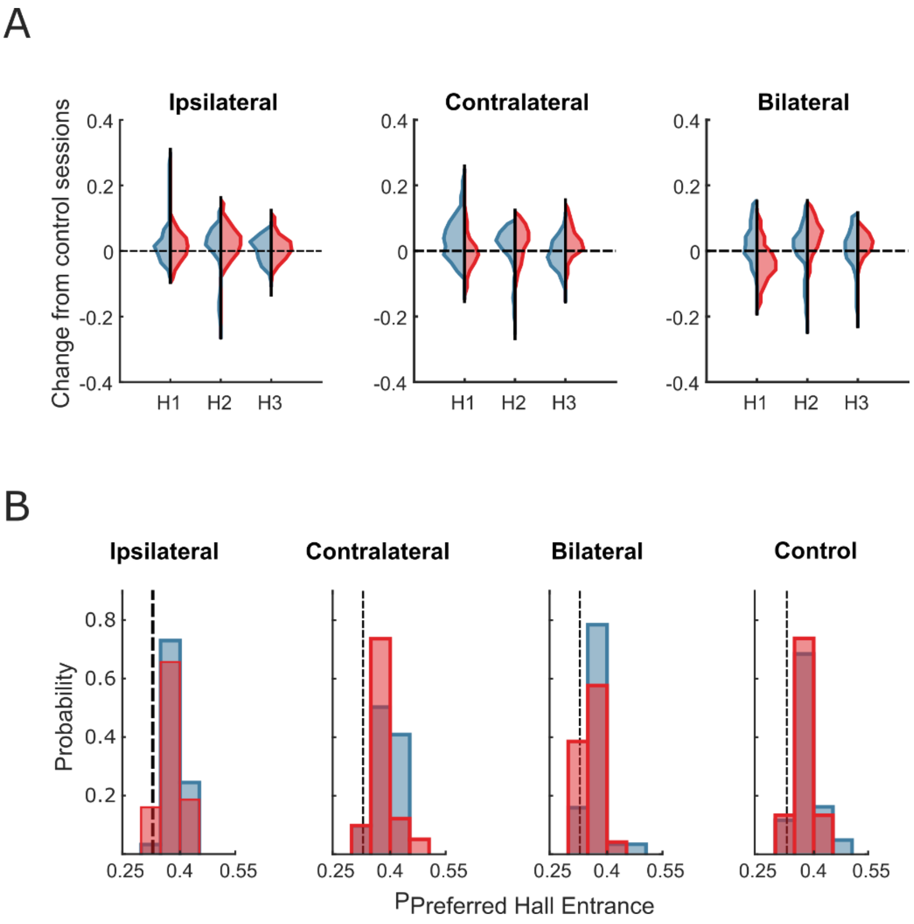


**Supplemental Figure 4: Stimulation did not lead to a conditioned hall preference**. **A)** Distributions of within-animal baseline-subtracted entrances into each hall. Stimulation of dSPNs (blue distributions) or iSPNs (red distributions) did not significantly alter the number of entrances to any hall regardless of which hemisphere(s) in which stimulation was presented (*p>0.05*, 2-way ANOVA with Bonferroni multiple comparison correction). The exceptions to this were the entrances into hall 1 and hall 3 during dSPN ipsilateral stimulation sessions (*p=0.0012*, 2-way ANOVA with Bonferroni multiple comparison correction) and contralateral stimulation sessions (*p=0.13*, 2-way ANOVA with Bonferroni multiple comparison correction). Given that the differences in hall entrances are not consistent between these two stimulation conditions and that no effect was seen as a result of iSPN stimulation, we argue that these differences in hall entrances do not represent conditioned hall preference. **B)** Probability of entering the most preferred hall in ipsilateral, contralateral, and bilateral stimulation sessions and control sessions. There was no significant difference between the probability of entering the most preferred hall in dSPN (blue) or iSPN (red) stimulation sessions or between stimulation sessions and control sessions (*p>0.05*, 2-way ANOVA with Bonferroni multiple comparison correction).
